# Fortuitous events in the evolution of Light-dependent Protochlorophyllide Oxidoreductase

**DOI:** 10.1101/2023.04.16.537069

**Authors:** Pratishtha Vedalankar, Baishnab C Tripathy

**Author notes:** Corresponding author Correspondence should be addressed to Professor Baishnab C Tripathy Department of Biotechnology Sharda University Greater Noida Uttar Pradesh 201310 India.

## Abstract

Light-dependent protochlorophyllide oxidoreductase (LPOR) is a nuclear-encoded photoenzyme in many photosynthetic organisms. LPOR originated in primitive cyanobacterial ancestors during the great oxygenation event that was detrimental to the existence of the oxygen-sensitive LIPOR that prevailed in anoxygenic Earth. Both LIPOR and LPOR catalyse reduction of protochlorophyllide to chlorophyllide in the penultimate step of chlorophyll biosynthesis. Except for angiosperms and gnetophytes several oxygenic phototrophs harbour both LIPOR and LPOR. The coexistence of LIPOR and LPOR in certain phototrophs provides niche spaces for organisms in unconducive environment. The selection pressure of increased O_2_ concentration, changing light quality and quantity at different depths of the ocean, nutrient status of water, gene reorganization during several endosymbiotic events, horizontal gene transfer, LIPOR gene loss and multiple duplication events played a major role in the evolution and diversification of LPOR and its isoforms in photosynthetic and non-photosynthetic organisms. In the absence of LIPOR angiosperms become vulnerable to protochlorophyllide-sensitized and light-induced oxidative stress mediated by singlet oxygen. To overcome the photo-damage PORA was expressed abundantly in the plastids of etiolated plants. PORB evolved to take over the function of vanishing PORA isoform in light. Brassicales evolved PORC to protect plants from high light and other environmental stresses.

**Highlights:**

1. Protochlorophyllide oxidoreductase is an important photo-enzyme in angiosperms that needs light as a substrate for the synthesis of chlorophylls. Therefore, angiosperms cannot green in dark although several algae and lower green plants can synthesize chlorophyll in dark due to the presence of light-independent protochlorophyllide oxidoreductase (LIPOR).
2. In response to climate change, during the great oxygenation event light-dependent protochlorophyllide oxidoreductase (LPOR) evolved due to the O_2_-induced selection pressure that inactivated the oxygen-sensitive LIPOR.
3. Increased O_2_ concentration, changing light quality and quantity at different depths of ocean, gene reorganization during several endosymbiotic events, selective LIPOR gene loss and multiple duplication events played a major role in the evolution and diversification of LPOR and its isoforms in phototrophs.
4. Phylogenetic studies indicate that LPOR genes have been overwhelmingly horizontally transferred between phototrophs and also non phototrophic organisms.
5. Presence of LPOR in non-photosynthetic organisms, Mycobacterium and certain fungi suggests that LPOR may have some other reductive functions in these organisms.

## 1. Introduction

Photosynthesis involves the conversion of the solar energy into chemical energy using readily available raw materials. What makes photosynthesis truly exceptional is that not only does it synthesize the building blocks of life, but it also provides the critically important oxygen for the much-needed survival of higher forms of living organisms (Björn, 2009; Blankenship, 2010). The status quo of photosynthetic processes was not always as we see it now. Evolutionary processes have left an indelible imprint on the timeline of photosynthesis. When the first photosynthetic organisms originated around 3.4 billion years ago (bya), earth was dominated by anoxygenic phototrophs that used hydrogen sulphide or other substrates as electron donors without the evolution of oxygen. The prokaryotic oxygenic phototrophs that originated in anoxygenic earth created the great oxygenation event (GOE) approx. 2.2 to 2.7 bya (Björn, 2009; Blankenship, 2010; Hohmann-Marriott and Blankenship, 2011; Buick, 2008; Schirrmeister et al., 2013; Fujita and Uesaka, 2022). This established the oxygenic photosynthesis on earth. The primary endosymbiotic event that gave rise to the classical double membrane bound organelles chloroplast and mitochondria occurred around 1.5 bya. Endosymbiosis culminated into the evolution of present day eukaryotic oxygenic photosynthetic organisms that use light energy for the oxidation of water, thereby releasing oxygen (Blankenship and Hartman, 1998; Archibald 2009;2015; Blankenship, 2010; Rebeiz et al., 2010; Hohmann-Marriott and Blankenship, 2011; Cardona, 2019).

Photosynthesis takes place with the help of the tetrapyrrolic light absorbing pigment molecules bacteriochlorophylls (BChls) and chlorophylls (Chls) in anoxygenic and oxygenic phototrophs respectively (Chen et al. 2010; Nascimento et al. 2016). The light-dependent reaction takes place in photosynthetic membranes and begins with photon absorption by pigment molecules associated with antennae proteins, followed by excitation energy transfer to the reaction centres (RC). The absorbed solar energy is conserved in the form of high-energy bonds of adenosine triphosphate (ATP), and a strong reductant, the reduced nicotinamide adenine dinucleotide phosphate (NADPH) (Rabinowitch, 1965; Rabinowitch and Govindjee, 1969; Bryant et al., 2020). There are many different types of Chls (Chla, Chlb, Chlc, Chld, Chle, Chlf, Chlg) and Bchls (Bchl a, Bchl b, Bchl c, Bchl d, Bchl e, Bchl f, Bchl g) (Chen et al., 2010; Kobayashi et al., 1998; Fujita and Yamakawa, 2017; Bryant et al., 2020). In oxygenic phototrophs Chl a is the most abundant Chl species which is a product of several enzymatic steps (depicted in Fig 1.) some of which are common for the biosynthesis of Bchl and Chl molecules (Chen et al., 2010; Scheer, 2006, Tripathy and Pattanayak, 2012).

**Figure 1.**
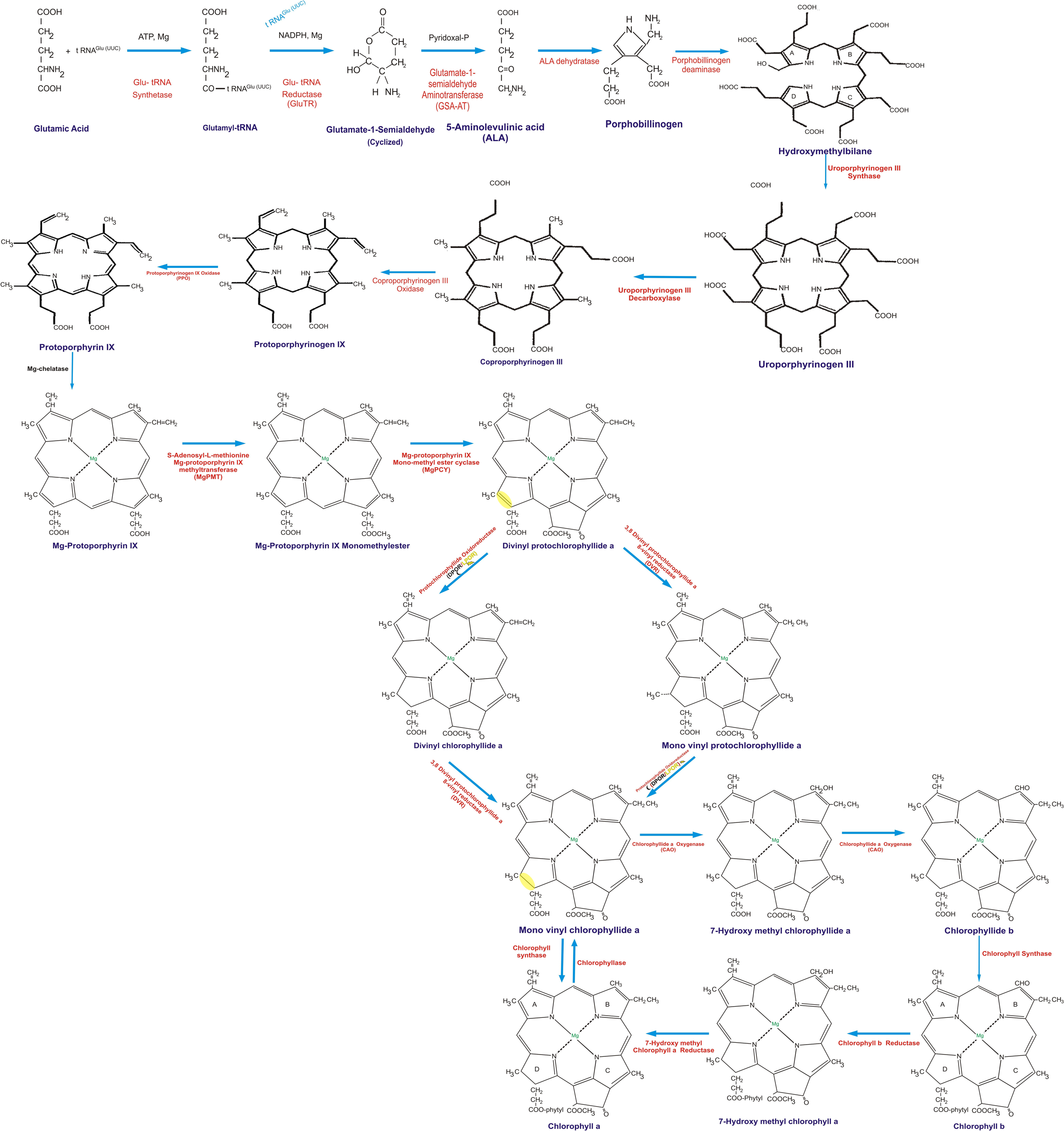
The schematic representation of the enzymatic steps involved in chlorophyll biosynthesis pathway starting from Glutamic acid

5-Aminolevulinic acid (ALA) is the first committed precursor for the synthesis of Mg-tetrapyrroles such as the Chls, Fe-tetrapyrroles, heme, cobalamin (vitamin B12), siroheme, and coenzyme F430 (Battersby, 2000; Hunter and Ferreira, 2009). In several bacteria and higher phototrophs ALA is synthesized from glutamyl-tRNAGlu by the activities of two distinct enzymes glutamyl-tRNA reductase (GluTR) and glutamate 1-semialdehyde aminotransferase (GSA-AT). However, ALA is synthesised from succinyl CoA and glycine in β proteobacteria, fungi and animals catalysed by ALA synthase. The biosynthesis of Chls, Heme or Bilin is similar till the formation of non-conjugated macrocyclic tetrapyrrole protoporphyrin IX from ALA (Beale, 1999; Tripathy and Dalal, 2013). At the onset condensation of two ALA molecules results in the synthesis of a 5-membered heterocyclic ring of porphobilinogen. Thereafter four molecules of porphobilinogen assemble to form the linear tetrapyrrole hydroxymethylbilane (Tanaka and Tanaka, 2007; Tripathy and Pattanayak, 2012; Bryant et al., 2020). Cyclization of hydroxymethylbilane forms uroporphyrinogen III with the inversion of the pyrrole D ring (Tanaka and Tanaka, 2007; Tripathy and Pattanayak, 2012; Brzezowski et al., 2015). Uroporphyrinogen-III is decarboxylated to produce coproporphyrinogen III. In addition to the synthesis of Chl, uroporphyrinogen-III serves as the substrate for siroheme biosynthesis. The two propionate side chains on the rings A and B of coproporphyrinogen III are oxidatively decarboxylated to produce protoporphyrinogen IX (Bollivar, 2006; Rebeiz et al., 2010). With the removal of 6 electrons from the macrocycle of protoporphyrinogen IX, aromaticity is conferred to the macrocycle to generate protoporphyrin IX. Protoporphyrin IX is the branch point for heme and Chl biosynthesis. The iron branch begins with the insertion of Fe^2+^ to protoporphyrin IX for heme biosynthesis (Tanaka and Tanaka, 2006; Scheer, 2006; Granick and Beale, 1978; Castelfranco and Beale, 1983; Carey et al., 1985). The insertion of Mg^2+^ ion into protoporphyrin IX converts it into Mg-protoporphyrin IX. Mg-protoporphyrin IX is esterified to form Mg protoporphyrin IX monomethyl ester that is subsequently metabolised by addition of a fifth isocyclic ring to synthesise Divinyl Protochlorophyllide (DV Pchlide). Protochlorophyllide (Pchlide) is the last common intermediate to a vast array of Chls (including Chl a, Chl b and BChls). It is also the bifurcation point for the formation of the Chl c family. In many phototrophs DV Pchlide is often acted upon by vinyl reductase to form Monovinyl protochlorophyllide (MV Pchlide) (Monovinyl plants) mostly during night time. MV Pchlide and DV Pchlide are acted upon by one of the key enzymes of Chl biosynthesis pathway, protochlorophyllide oxidoreductase (POR) that catalyses the reduction of C17 – C18 double bond of D ring of Pchlide to synthesise Divinyl Chlorophyllide a (DV Chlide *a*) and Monovinyl chlorophyllide *a* (MV Chlide *a*) respectively. DV Chlide *a* so formed is swiftly converted to MV Chlide *a* by vinyl reductase (Duggan and Rebeiz, 1982; Tripathy and Rebeiz, 1986; 1988; Tripathy and Pattanayak, 2012). Chlorophyllide *a* (Chlide *a*) is metabolised to chlorophyllide *b* (Chlide *b*) by chlorophyllide a oxygenase (CAO) and both Chlide *a* and Chlide *b* are esterified with phytol to form Chl a or Chl b (Duggan and Rebeiz, 1982; Tripathy and Rebeiz, 1986; 1988).

## 2. Protochlorophyllide oxidoreductase

POR exists in two different non-homologous enzymatic forms in phototrophs-(1) NADPH Light dependent Protochlorophyllide Oxidoreductase (LPOR) and (2) Light Independent or Dark Operative Protochlorophyllide Oxidoreductase (DPOR/LIPOR) (Griffiths, 1978; Fujita, 1996; Adamson et al., 1997; Brzezowski et al., 2015; Vedalankar and Tripathy, 2019). Even though both the POR enzymes catalyse the same reaction they differ from each other right from their origin, subunit composition, structure, catalytic mechanism, and their spread and diversity across the photosynthetic organisms (Hunsperger et al., 2015).

LIPOR originated under anoxygenic conditions in the reducing atmosphere of primitive earth. It is widely distributed among photosynthetic organisms ancient anoxygenic to oxygenic phototrophs including cyanobacteria, algae, bryophytes, pteridophytes and gymnosperms except gnetophytes and angiosperms (Bauer et al., 1993; Suzuki and Bauer, 1995; Fujita, 1996; Boivin et al., 1996; Armstrong, 1998; Walmsley et al., 1999; Breznenová et al., 2010; Reinbothe et al., 2010). The reduction of Pchlide by LIPOR takes place in the absence of light in an ATP dependent reaction using ferredoxin as reductant (Bröcker et al., 2008; Nomata et al., 2016). LIPOR is encoded by three genes (BchL, BchN, BchB/ChlL, ChlN, ChlB) (Burke et al., 1993a; 1993b, Suzuki and Bauer, 1992; Bröcker et al., 2008; Fujita, 1996). Structurally LIPOR is homologous to the ancient Nitrogenase enzyme and consists of two separable protein components: the L protein and the NB protein that are analogous to the Fe-protein and MoFe protein complex of the nitrogenase (Armstrong, 1998; Fujita and Bauer, 2000). The reductive L protein complex is a homodimer of bchLbchL/chlLchlL subunits, and the catalytic NB protein complex is composed of a heterotetramer of 2bchNbchB/2chlNchlB subunits (Fujita and Bauer, 2000; Bröcker et al., 2010).

In contrast to LIPOR that is coded by the chloroplast genome, LPOR is a nuclear encoded, single polypeptide of approx. 36kda and has an absolute requirement of light for catalysing the reduction of Pchlide to Chlide (Mullet, 1988; Hunsperger et al., 2015). It requires NADPH as a cofactor and having evolved in the oxygenic environment it is prevalent in all oxygenic phototrophs; unlike LIPOR, the LPOR is insensitive to oxygen (Armstrong, 1998; Bollivar, 2006; Yamamoto et al., 2009). LIPOR is believed to have originated prior to GOE and hence it is considered to be the ancient Pchlide reducing enzyme (Yang and Cheng, 2004; Björn, 2009; Schirrmeister et al., 2013). It is widely believed that the GOE triggered the evolution of LPOR during earth’s transition from reducing to oxidising atmosphere created by O_2_ evolving cyanobacteria that converted oceanic water to oxygen (Buick, 2008). Evolutionarily both the Pchlide reducing enzymes, LIPOR and LPOR coexist in many photosynthetic organisms (Yang and Cheng, 2004; Yamazaki et al., 2006; Vedalankar and Tripathy, 2019). This review discusses the conundrum of origin and evolution of LPOR, a key light-driven enzyme that plays a crucial role in the Chl biosynthesis and plant development. A better understanding of the LPOR would allow us to understand the need for the existence of entirely two different enzymatic processes for Pchlide reduction i.e., the selection pressure that upended the light-independent reduction to a completely different light-dependent process (Suzuki and Bauer, 1995).

It has been well established that light is an indispensable component for the activity of LPOR enzyme much like the DNA repair enzyme DNA photolyase (Begley, 1994; Björn, 2018), bacterial chlorophyllide a reductase (COR) (Saphier et al., 2005), cyanobacterial chlorophyllide f synthase (Chen et al., 2010; Galperin et al., 1998; Ho et al., 2016) and fatty acid photodecarboxylase (FAP) (Sorigué et al., 2017). In addition to light, LPOR requires NADPH as a reductant and Pchlide as a target substrate to catalyse the stereospecific reduction of the C17-C18 double bond in the porphyrin D ring of (Pchlide a) -to (Childe-a) (Yang and Cheng, 2004; Gabruk and Mysliwa-Kurdziel, 2015). Photoreduction of Pchlide to Chlide is an ultrafast event that involves transient charge separation across the C17-C18 double bond of the pigment leading to the formation of charge transfer intermediates which facilitate the step wise hydride and proton transfer. These intermediates have been analysed on an ultra-fast time scale by time resolved fluorescent measurements. The proton transfer to C18 of Pchlide by charge transfer intermediates occurs at approx. 3 picoseconds during the first dark reaction while the rate limiting hydride transfer step from pro-S face of NADPH to C17 of Pchlide occurs at 400 ps. The rate of hydride transfer in Pchlide photoreduction is faster in eukaryotes as compared to prokaryotes suggesting that efficient LPOR evolved during the endosymbiotic era (Heyes and Hunter, 2005; Heyes et al., 2006;2021).

## 3. Epochal events leading to the origin of LPOR

The redox state of the earth’s atmosphere changed dramatically around 2.45 Ga years making free O_2_ a permanent constituent of the earth’s atmosphere mostly due to the recruitment of Mn-containing oxygen-evolving complex proteins and the evolution of chlorophyll by cyanobacteria that utilized the absorbed solar energy for photolysis of water (Van Kranendonk et al., 2012). This oxygenic photosynthesis allowed the evolutionary emergence of eukaryotes, multicellular organisms, and complex life forms as we know them. The cyanobacterial diversity indicates the origin of cyanobacterial ancestors sometime in the Archean era. The rise of oxygen during the GOE exerted evolutionary pressure on the 4Fe-4S center containing the oxygen-sensitive L subunit of LIPOR and triggered the origin of the oxygen-insensitive Pchlide-reducing enzyme LPOR (Olson, 2001; Schoefs and Franck, 2003; Yamazaki et al., 2006; Nomata et al., 2006; Yamamoto et al., 2009; Reinbothe et al., 2010; Hunsperger et al., 2015). It is noteworthy that the oxygen level present during the late Proterozoic era just after GOE was adequate for the evolution of LPOR (Olson, 2001; Björn, 2009; Blankenship, 2010; Schirrmeister et al., 2013; Shih et al., 2013). Similarly, cyanobacteria having another O_2_-sensitive enzyme nitrogenase, like LIPOR, having a monopolistic dominance in the anoxygenic environment (Fujita and Bauer, 2000) acquired temporal differentiation by carrying out nitrogen fixation in the dark and spatial differentiation by confining nitrogen fixation to specialized cells called heterocysts. These heterocystous cyanobacteria evolved late in the O_2_ event to protect themselves from the oxygenic threat (Fujita and Bauer, 2000; Yamazaki et al., 2006; Fujita and Uesaka, 2022). However, a similar mechanism was not observed in LIPOR, and it is speculated that LIPOR acquired some other protective mechanisms such as the water-water cycle to remove oxygen to combat the oxygen sensitivity (Durnford and Falkowski, 1997). The evolution of LPOR did not lead to the extinction of LIPOR. Both the nonhomologous Pchlide-reducing enzymes coexist among the oxygenic phototrophs, the only exceptions being gnetophytes and angiosperms where LIPOR is non-existent (Walmsley et al., 1999; Schoefs and Franck, 2003; Yamazaki et al., 2006; Hunsperger et al., 2015). Why the two enzymes still coexist today even in advanced archegoniate like gymnosperms does not have a precise answer.

Evolution and spread of LPOR took a magnificent jump to higher eukaryotes that acquired photosynthesis via endosymbiosis. The primitive cyanobacterium was established as a photosynthetic organelle called plastid inside the eukaryotic phototrophs about 1.5 bya (Archibald, 2009; Chan and Bhattacharya, 2010; Keeling, 2010).The photosynthetic eukaryotic ancestor gave rise to the complex supergroup Archiplastida that branched out into three separate photosynthetic lineages – the Glaucophyta, Viridiplantae (comprising green algae and land plants), and Rhodophyta –the red algae. Subsequently, the secondary, tertiary, and higher-order endosymbiosis transmitted the plastids amongst eukaryotes to give rise to the present-day diversity of eukaryotic photosynthetic lineages (Tomitani et al., 1999; Chan et al., 2010; Keeling, 2010; Archibald and Keeling, 2002; Archibald 2015). The endosymbiotic process has played a crucial role in the evolution of photosynthetic genes in eukaryotes. A major aftermath of endosymbiosis was a massive transfer of genes from the endosymbiont to the host nucleus (Keeling, 2010). The endosymbiotic genes underwent several recombination events and became integrated into the host nuclear genome. As a result, the size of the plastid genome was significantly reduced compared with the original cyanobacterial endosymbiont. After primary endosymbiosis, the LPOR genes were transferred to the nucleus, while the LIPOR genes remained in the chloroplast genome or were lost in several algal lineages and higher plants in the oxygenic environment (Hunsperger et al., 2015). Until recently only a single endosymbiotic gene transfer event was thought to have occurred that led to the origin of plastids by engulfment of the β-Cyanobacteria possessing β-carboxysomes with Form-IB Rubisco (Kerfeld and Melnicki, 2016). About 60 million years ago another independent endosymbiotic event took place in freshwater rhizarian amoeba *Paulinella chromatophora* (Nowack, 2014; Kim and Park, 2016). These organisms retain cyanobacterial prey of the Prochlorococcus and Synechococcus sp PCC 7002 of α Cyanobacteria type which has α-carboxysomes with Form-IA Rubisco (Kerfeld and Melnicki, 2016; Gabr et al., 2020). The endosymbiotic photosynthetic entities in *P. chromatophora* are called as chromatophores that absorb solar energy and they retain cyanobacterial features such as carboxysomes, phycobilisomes, peptidoglycan cell wall (Nowack and Grossman, 2012; Kerfeld and Melnicki, 2016; Kim and Park, 2016). This is a chloroplast in making.

In prokaryotes, horizontal gene transfer (HGT) is ingrained within the genealogical fabric of the organisms (Treangen and Rocha, 2011; Bock, 2010; Koonin, 2016). The numerous instances of HGT in organisms although sporadic give ample evidence of the important and pervasive role of lateral gene transfer (Bock, 2010; Brinkmann et al., 2018). The temporal intrusions in HGT are a result of a unique combination of begging or borrowing or stealing genetic information across the natural mating barriers (Ku and Martin, 2016; Bock, 2010). The presence of ancestral, novel genes or paralogs of existing genes or xenologous sequences can be estimated from the phylogenetic analysis depicting the divergence of the recipient from the direct ancestor (Ku and Martin, 2016; Cohen et al., 2011; Hunsperger et al., 2015). The HGT has contributed immensely towards the evolutionary process of LPOR in photosynthetic organisms. LPOR phylogeny follows the identical pattern to the plastid transfer among algae originating from primary endosymbiosis (Hunsperger et al., 2015). Typically, plastids derived from green algae fall into three separate lineages streptophytes, chlorophytes, and primitive prasinophytes (Archibald and Keeling, 2002). The phylogenetic studies reveal that the marine phototrophic amoeboid flagellates chlorarachniophytes evolved from the chlorophyte clade (Chlorophyceae– Trebouxiophyceae– Ulvophyceae-Pedinophyceae) of green algae (Cavalier-Smith, 1998). The chlorarachniophyte plastids typically contain green algal footprints but some members of chlorarachniophyte including *Bigelowiella natans* contain plastid proteomes and nuclear genes heavily infected with LPOR genes originating from phylogenetically divergent red algae or from xenologous sources (Curtis et al., 2012; Hunsperger et al., 2015). The abundance of ‘foreign’ genes in *Bigelowiella natans* as a result of HGT compensates for the mixotrophic lifestyle (Rogers et al., 2007; Curtis et al., 2012; Burki, 2017). LPOR proteins originating from the green algae lineage due to the secondary endosymbiotic event are found to be sister branches to one another and nested within the prasinophyte algae indicating a possibility of HGT from prasinophyte to the chlorarachniophyte (Hunsperger et al., 2015). The LPOR proteins funneled from the Rhodophyta branch of plastids should belong within the rhodophytic clade but there are numerous instances with most members following a rather reticulate path similar to that observed due to secondary and higher-order symbiosis. The exact origins of the LPOR genes transferred via red algal lineage are complex (Hunsperger et al., 2015). Phylogenetic studies demonstrate that maximum members of stramenopiles demonstrate an affinity to the prasinophytes branch. All sequenced haptophytes and quite a few peridinin-containing dinoflagellates show their origin within the prasinophytic branch of LPOR proteins (Hackett et al., 2004; Minge et al., 2010). Either the primary Rhodophyta LPOR genes have been completely replaced by HGT to the chlorophyte genes. Within the Cryptophytes and Heterokonts bearing rhodophytic LPORs several phylogenetic reconstructions are observed originating from EGT and several independent HGTs (Keeling, 2010; Hunsperger et al., 2015). Diatoms acquired plastids via secondary endosymbiosis from the red algal lineage (Armbrust et al., 2004; Janouškovec et al., 2010). The diatom *Alexandrium tamarense* shows presence of prasinophytic LPOR genes acquired from green alga *Micromonas* sp as a result of HGT (Wisecaver et al., 2013; Hunsperger et al., 2015). The recently sequenced genome of a number of diatoms reveals a chimeric or transgenic genome that shows several traces of green algae footprints and xenologues acquired from a variety of other sources by EGT and HGT (Petersen et al., 2014). This feature enables diatoms to survive in difficult habitats and varying environmental conditions.

## 4. Migration of LPOR to anoxygenic photosynthetic organisms

Like millions of oxygenic photosynthetic species possess LIPOR, several aerobic anoxygenic phototrophic bacteria (AAPB) have functional LPOR sequences. AAPBs are a ubiquitous group of marine microbes that possess light harvesting rection centers supplemented with heterotrophic metabolism. a globally conserved structure similar to the well-characterized cyanobacterial LPOR (Biebl et al., 2005; Chernomor et al., 2021). In contrast to anoxygenic photosynthetic bacteria (APB) like *Rhodobacter capsulatus* AAPBs can perform anoxygenic photosynthesis in the presence of atmospheric oxygen (Yurkov and Hughes, 2017). Possession of LPOR by AAPBs is an adaption to increase Bchl synthesis under aerobic conditions. Members of α proteobacteria - *Dinoroseobacter shibae*, *Erythrobacter litoralis*, *Yoonia vestfoldensis*, *Sulfitobacter guttiformis*, *Porphyrobacter dokdonensis* and β proteobacteria – *Limnohabitans* sp. and *Burkholderia* sp acquired LPOR probably due to HGT from cyanobacteria like organisms involving multiple gene transfer events that might have taken place at different time points (Biebl et al., 2005; Brinkhoff et al., 2010; Wagner-Döbler et al., 2010;Kaschner et al., 2014;Wang et al., 2014; Chernomor et al., 2021; Bryant and Frigaard, 2006; Yurkov and Hughes, 2017; Kasalický et al., 2018). Anoxygenic phototroph *Gemmatimonas phototrophica* member of phylum Gemmatimonadetes also acquired LPOR as an inter-phylum HGT from proteobacteria (Zeng et al., 2015; Chernomor et al., 2021). Phylogenetic studies by Chernomor et al. 2021 expand the database of LPOR sequences in AABPs. The refuge of LPOR in AAPBs is part of a design rather than a chance event or an accidental event which turns favourable and this is not restricted to a few proteobacterial species thus firmly establishing the presence of LPOR in AAPBs (Chernomor et al., 2021).

## 5. Gene duplication

Gene duplication is another major mechanism that introduces genomic novelty among organisms (del Pozo and Ramirez-Parra, 2015). The duplication event not only increases the copy number of the duplicated gene but with time a copy of the duplicated gene might accumulate mutation and acquire a novel function in the genome (Birchler and Yang, 2022; Savino et al., 2022). In many cases, the duplicated gene ends up as a non-functional counterpart which does not make a functional product due to modifications in the DNA sequence by nucleotide insertions, deletions, frameshifts, or substitutions that disrupt the reading frame or lead to the insertion of a premature stop codon (Hunsperger et al., 2015; Savino et al., 2022). Three algae lineages underwent genome duplications resulting in LPOR duplication: dinoflagellates, chlorarachniophytes, haptophytes, and stramenopiles (Janouškovec et al., 2017).

The Brassicales underwent three whole genome duplications and the three isoforms of LPOR in *Arabidopsis* may be the outcome of these events (Oosawa et al., 2000; Pattanayak and Tripathy, 2002). LPOR expansion in different photosynthetic organisms shows that it exists in three different isoforms within the genome of photosynthetic organisms (Yang and Cheng, 2004). The LPOR genes in Brassicales are very similar at the nucleotide level which probably reflects recent gene duplication (Oosawa et al., 2000). LPOR gene duplication occurred after the primary endosymbiotic event in eukaryotic plant cells (Archibald, 2015). The LPOR isoforms in some photosynthetic eukaryotes share similar functions and regulations even though they have been shown to arise from unique duplication events during evolution (Hunsperger et al., 2015). The evidence for the gene duplication event comes from phylogenetic trees which show the distribution of each of the two gene copies between two principal branches of the tree (Hunsperger et al., 2015). Studies show that euglenids and chlorarachniophytes arose from two unique duplication events. *Hordeum vulgare* and *A. thaliana* LPOR sequences arose independently but they have similar functions in seedling greening and Chl synthesis.

Recent studies have identified multiple LPOR genes arising as a result of duplication in several diatom genomes including *Phaeodactylum tricornutum*, *Thalassiosira pseudonana* (Armbrust et al., 2004) *Fragilariopsis cylindrus*, and *Pseudo-nitzschia multiseries*. Two LPOR isoforms (POR1 and POR2) present in *Phaeodactylum tricornutum* genome are the result of gene duplication to enable it to adapt under varying light conditions (Ambrust et al., 2004; Hunsperger et al., 2015). Certain members of cryptophytes show LIPOR gene loss but other members retain functional LIPOR genes. In cryptophyte, *Chroomonas mesostigmatica* CCMP1168 some of the sequences of chlN occur as a pseudogene, stop codons, and frameshifts in the nuclear genome. The chlL is absent from *Rhodomonas salina*, *G. theta*, and *H. andersenii* plastid or nuclear genome (Fong and Archibald, 2008; Hunsperger et al., 2015; Kim et al., 2017). Dinoflagellates *Heterocapsa triquetra* (Fong and Archibald, 2008) and *Amphidinium* (Lauritano et al., 2017) show the presence of only one subunit of chlL which has shown to be a functional copy by expressed sequence tag surveys. LIPOR subunit in *H. andersenii* and *Chroomonas pauciplastida* possess group II introns in different locations making the gene non-functional (Khan and Archibald, 2008). *Arabidopsis thaliana* and *Oryza sativa* contain approx 2700 and 5600 well-defined pseudogenes, respectively (del Pozo and Ramirez-Parra, 2015). Pseudogenes are non-functional but still beneficial in evolution as it helps the organism to counter or survive the changed environmental conditions (Treangen and Rocha, 2011). Plant genomes are replete with instances of the presence of pseudogenes that are non-functional but still beneficial in evolution as it helps the organism to counter or survive the changed environmental conditions. Several algae that lack LIPOR, including certain haptophytes and stramenopiles, the LPOR gene is duplicated, and it is hypothesized that this duplication might compensate for the loss of the dark enzyme, potentially by allowing for the differential regulation of the genes of the photoenzyme (Hunsperger et al., 2015).

## 6. Impact of other Environmental factors on LPOR origin and evolution

Environmental factors often determine the distribution and diversification of genes for appropriate adaptation. Genes are rendered inactive by a mutation or gene loss and become dispensable during evolutionary pressure. Although this might not be a universally adaptive process, it is helpful under certain situations to combat the evolutionary pressure (Helsen et al., 2020). Pchlide reduction has endured various selective pressures across a range of environmental parameters after the transition of photosynthesis from an initial anoxygenic to an oxygenic environment. Some of the environmental factors include a) High altitude, b) Fe deficiency and c) high temperature, d) light intensity, e) light quality (Li and Bridwell-Rabb, 2018; Przybyla-Toscano et al., 2021; Abbas et al., 2022; Kees et al., 2022).

In the oxygenic environment, O_2_-dependent MPE (oxidative) cyclase is responsible for Pchlide synthesis (Figure 1). A decrease in the partial pressure of O_2_ reduces the Pchide concentration in plants (Li and Bridwell-Rabb, 2018). Therefore, the POR activity is downregulated and it influences the distribution and accumulation of the Pchlide and its reducing enzyme POR at different altitudinal clines (Abbas et al., 2022). In the early oxygenic world, the lower partial pressure of oxygen was sufficient to generate stratospheric O3 in the presence of UV light that prevented the UVB penetration and DNA damage of haploid life forms and produce ROS including singlet oxygen (^1^O_2_). ROS was not only responsible for the destruction of Fe-S centers of LIPOR in several species it also promoted the evolution of different isoforms of LPOR. However, this O_2_ concentration was limiting for MPE (oxidative) cyclase, and therefore, it impacted Pchlide synthesis, LPOR evolution and Chl synthesis in early oxygenic haploid photoautotrophs.

Similarly, nutrient deficiency especially Fe impacted LPOR evolution with concomitant loss of LIPOR. The 4Fe-4S centers of L subunit of LIPOR needs Fe, for its synthesis and assembly. Oceanic diatoms Phaeodactylum tricornutum, Thalassiosira pseudonana and Nannochloropsis oceanica lost LIPOR because they are majorly present in Fe-sufficient oceanic water (Przybyla-Toscano et al., 2021). The low availability of micronutrient Fe encouraged the dispensability of LIPOR and the establishment of LPOR for better adaptability of other oceanic phototrophs that acquired LPOR by horizontal gene transfer (Behrenfeld et al., 2006; Bowler et al., 2010; Cvetkovska et al., 2019)

The LPOR activity in different species is also temperature dependent. In Synechocystis the optimum LPOR activity is at 30^0^C. However, in thermophilic cyanobacterium Thermosynechococcus elongates the optimum temperature for LPOR activity is between 50^0^C-55^0^C and it is less active at room temperature (McFarlane et al., 2005). Although cyanobacteria mostly possess both the Pchlide reducing enzymes i.e., LIPOR and LPOR; thermophilic cyanobacteria lack both LIPOR genes and nitrogenase genes. This loss of LIPOR genes is attributed to high temperature-induced genome reduction in thermophilic cyanobacteria (Kees et al., 2022).

Light intensity is a factor influencing the light-driven reaction rates. The LPOR disrupted YFP12 mutant of cyanobacteria and wild-type cyanobacteria grow normally under low light conditions (10-25 mu E m^-2^ s^-1^), but at high light intensity (85-170 mu E m^-2^ s^-1^), the mutants stop growing and are photo-bleached (Fujita et al., 1998). In contrast YFC2 mutant with disrupted ChlL (LIPOR less) grew rapidly at higher light intensities, suggesting that the contribution of LPOR in chlorophyll biosynthesis increases with increase in light intensity (Fujita et al., 1998). Under medium-light intensities (25–130 mu E m^-2^ s^-1^) as well as high-light intensities (above 130 mu E m^-2^ s^-1^) LPOR is majorly and exclusively functional for Chl biosynthesis (Fujita et al., 1998). LIPOR is more energy-demanding than LPOR and thus has been replaced during the evolutionary process to allow photosynthetic organisms to follow a more conservative lifestyle (Masuda and Takamiya, 2004). The contribution of LPOR towards the Chl biosynthesis increases with increasing light intensity. LPOR acts as a trigger for the germination of seedlings in plants and provokes a marked change in the morphological development of the plant (Ha et al., 2017). At low light intensities, Chl content in cyanobacterium Plectonema boryanum lacking LIPOR was lower, and its growth rate is retarded (Huang et al., 2004). The early cyanobacteria that had only LIPOR most likely adapted to low light intensity for efficient Chl biosynthesis and photosynthesis for survival. In this context, *Chlamydomonas* sp. UWO241 is an intriguing chlorophyceae member as despite inhabiting low light conditions for extended periods it shows loss of LIPOR, as a result, Pchlide reduction is entirely carried out by LPOR (Bowler et al., 2010; Smith et al., 2019). *Chlamydomonas* sp. UWO241 with its reduced chloroplastic genome inhabits perennially ice-covered lakes of Antarctica with high dissolved oxygen content, iron limiting, and hypersaline conditions. The above conditions seem to support the loss of LIPOR (Behrenfeld et al., 2006; Bowler et al., 2010; Cvetkovska et al., 2019; Smith et al., 2019). It seems that the ancestral chlamydomonadalean clade that gave rise to UWO241 seems to have lost the LIPOR genes (Cvetkovska et al., 2019; Smith et al., 2019).

The spectral composition of the light changes with topographic factors. Red light which is of longer wavelength is mostly absorbed by the water at the surface. Blue light penetrates deep and turbid water. The red light is highly efficient for the LPOR-mediated photoconversion of Pchlide to Chlide (Dalal and Tripathy, 2012). Nevertheless, the supposed inefficiency of LPOR in deep or turbid water is not surprising as in these conditions only blue light is available due to the scattering. Greenlight that is available deep in the ocean is not efficiently absorbed by many phototrophs (Kehoe and Gutu, 2006). The action spectra of LPOR reveals that it is ineffective in green light, in such cases LIPOR compensates for the low LPOR activity. The members of the (Microchaetaceae) *Fremyella diplosiphon* (Shui et al., 2009; Pattanaik et al., 2011), (Nostocaceae) *N. punctiforme* (Hirose et al., 2013), (Rivularariaceae) *Gloeotrichia 583* (Stowe et al., 2011) depict an interesting phenotype as an example of the complementary chromatic adaptation by alternating their photosynthetic pigments and enzymes under fluctuating light conditions (Hirose et al., 2013). It would not be hyperbolic to say that they can sense a rainbow of colors ranging from red, blue, green and violet (Grossman, 2003; Bordowitz and Montgomery, 2008; Kehoe, 2010). All these explain the simultaneous existence of LPOR and LIPOR for sustenance and adaptation to environmental factors.

### 6.1. Photoprotective role

The catalytic rate of LIPOR for the dark conversion of Pchlide to Chlide is quite slow. Therefore, under the steady-state conditions of Chl biosynthesis the Pchlide accumulation in the cells of LIPOR-containing organisms is high (Soffe, 2016). Under high light that prevails on the ocean surface, the accumulated Pchlide in LIPOR-containing organisms absorb light and transfer their energy to oxygen to produce highly reactive singlet oxygen (^1^O_2_) (Chakraborty and Tripathy, 1992; Tripathy and Pattanayak 2010; Pattanayak and Tripathy, 2011) that causes photooxidative damage to cells. The LPOR bestows photo-protection on the plants by limiting the Pchlide-mediated photo-oxidative damage (Buhr et al., 2008; Tripathy and Pattanayak, 2010, Pattanayak and Tripathy, 2011). Whereas the high light intensity on the surface of the ocean could photodamage the slow LIPOR-containing photoautotrophs, it can cause minimal damage to organisms possessing LPOR that converts Pchlide to Chlide rapidly within 1 millisecond (Sytina et al., 2008; Soffe, 2016; Heyes et al., 2021). Thus, LPOR protects the etiolated and green phototrophs by binding to the photosensitive Pchlide pool to keep it in photo-transformable form for very fast photo-conversion of Pchlide to Chlide to minimize generation of singlet oxygen that causes destruction of photosynthetic organisms in high light (Tripathy and Chakraborty, 1991, Chakraborty and Tripathy, 1992, Tripathy and Pattanayak, 2011). In angiosperms during senescence the chlorophyll content rapidly declines, however, POR activity persists to photo-transform protochlorophyllide to minimize singlet oxygen production to ensure leaf survival and the translocation of photosynthates from the source to sink (Hukmani and Tripathy, 1994).

The free Chls do not accumulate rather they bind to photosynthetic pigment-protein complex to efficiently transfer all of its absorbed energy to the reaction center for utilization. The antenna Chl molecules usually do not generate singlet oxygen (^1^O_2_) unless they are under high excitation pressure (Tripathy and Pattanayak, 2010; Pattanayak and Tripathy, 2011; Buhr et al., 2008). Therefore, high light acted as a selection pressure for the evolution of LPOR to protect photoautotrophs from photo-oxidative damage.

The LPOR genes are more widespread among phototrophic taxa and therefore, it is fair to interpret that LPOR genes are superior to LIPOR enabling plants to survive and grow at different environmental conditions. LIPOR genes often become redundant, and they are discarded under some environmental circumstances /selection pressure. Gene dispensability is often inferred by the higher rate of non-synonymous substitution rates and high ratio of non-synonymous to synonymous substitution. It has been shown previously that ChlL gene of the gymnosperm *Thuja standishii* has high rate of non-synonymous substitution confirming its dispensability (Kusumi et al., 2006). Thus, greater selection pressure acted on LIPOR genes leading to its complete absence from higher gymnosperm, gnetophytes, and angiosperms (Walmsley et al., 1999; Yamamoto et al., 2017). Taken together these findings suggest that LPOR genes seem to be to have higher functionality and importance compared to LIPOR genes in the oxygenic world.

## 7. Structural basis of LPOR function

### 7.1 LPOR a short-chain dehydrogenases/reductases superfamily confrère

LPOR belongs to a large family of enzymes known as short-chain dehydrogenases/reductases (SDRs) (Yang and Cheng, 2004; Wilks and Timko, 1995; Moummou et al., 2012). SDR is part of a large superfamily of enzymes known as the ‘RED’ (Reductases, Epimerases, Dehydrogenases) that catalyze a variety of NADP (H) - or NAD(P)+-dependent reactions (Wilks and Timko, 1995; Oppermann et al., 2003; Moummou et al., 2012) involving hydride and proton transfer (Hoeven et al., 2016; Archipowa et al., 2018). This is one of the oldest and most diverse protein families present in prokaryotes and eukaryotes that typically occur as oligomers (Oppermann et al., 2003; Yang and Cheng, 2004). It has a wide range of substrates involved in secondary metabolic routes ranging from polyols, retinoids, sterols, sugars, aromatic compounds, and xenobiotics (Persson et al., 2003). Plant LPORs are assigned to SDR73C family in the SDR superfamily (Dong et al., 2020).

The classical SDR family of proteins containing all oxidoreductases has two domains, one for binding of the cofactor and another for binding the substrate (Moummou et al., 2012). Despite the considerably low sequence similarity (15% -30%), SDR family members bear significant structural similarity such as a common a/ß folding pattern with Rossmann fold - a characteristic of SDR family members (Yang and Cheng, 2004). The Rossmann fold consists of a central parallel 6-7 β-sheet sandwiched by two arrays of 2-3 α-helices on either side making up αβα-core for NADPH binding (Dong et al., 2020). The Rossmann-fold and a hydrophobic loop region both are involved in anchoring the enzyme to the membrane (Moummou et al., 2012). SDRs consist of a one-domain subunit of about 250 amino acids with the cofactor binding site in the N-terminal part and substrate binding in the C-terminal part. The variable C-terminal segment determines the substrate specificity (Dong et al., 2020).

Other common structural features include a highly conserved active site with a highly conserved pentapeptide YxxxK in the catalytic motif (YKDSK in LPOR) that participates in the proper coordination with NADPH and Pchlide binding (Lebedev et al., 2001; Gabruk et al., 2016). The N terminal contains the conserved sequence (Gly-X-X-X-Gly-X-Gly) in SDR and GASSGV/LG in all LPORs. This glycine-rich motif is for structural integrity and binding of the pyrophosphate portion with NADPH (Dong et al., 2020). A key feature of the SDR superfamily is its catalytically important tetrad Ser-Asn-Tyr-Lys for proton transfer and stabilization of reaction intermediates. The catalytic triad in (Asn-Ser-Thr-Lys) POR contains Thr 145 instead of Ser residue (Moummou et al., 2012; Dong et al., 2020). Site-directed mutagenesis and in vivo analysis confirm that Tyr and Lys are the most conserved at the catalytic site in all LPOR members and that these are indispensable for the enzymatic catalytic activity (Wilks and Timko, 1995; Suzuki and Bauer, 1995; Lebedev et al., 2001 Heyes and Hunter, 2002). The Tyr residue acts as a general acid upon deprotonation and facilitates hydride transfer to or from NAD (P)+/H (Ehrig et al., 1994; Lebedev et al., 2001). The proton at the C-18 position of Pchlide is derived from Tyr and the hydride transferred to the C-17 position is derived from the pro-S face of NADPH (Heyes and Hunter, 2005; Archipowa et al., 2018). The mutation of either Tyr 275 or Lys-279 does not completely abolish the catalytic activity of LPOR. However, mutation of either residue impairs the formation of the ground state ternary enzyme-substrate complex, indicating their key role in substrate binding (Dahlin et al., 1999; Heyes and Hunter, 2002). Both residues have multiple roles in catalysis, involving formation of the ground state ternary enzyme-substrate complex, stabilization of a Pchlide excited state species and proton transfer to the reaction intermediate formed after the light reaction (Menon et al., 2009; Dong et al., 2020) (Figure 2).

**Figure 2.**
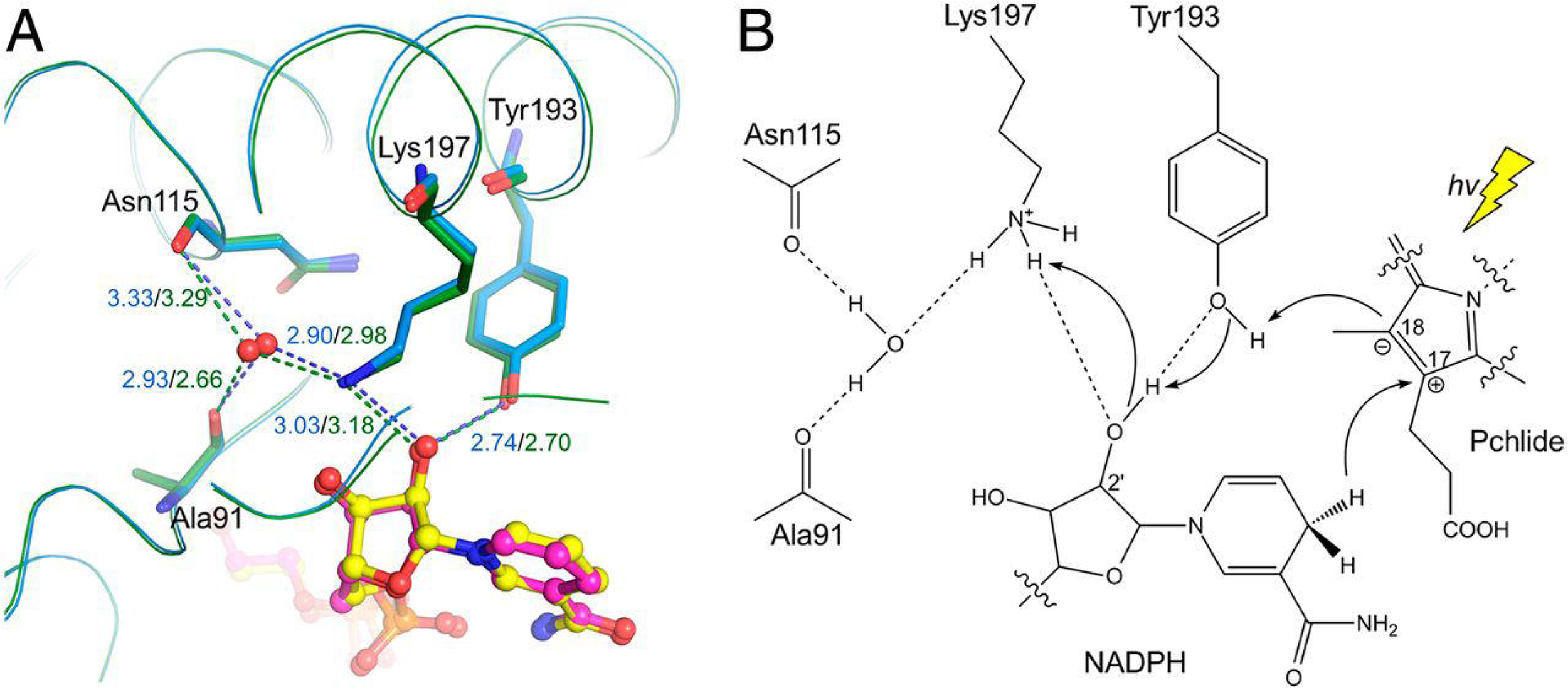
Proposed proton-relay path from Dong et al. 2020. (A) The hydrogen bond network bridging the Tyr193 ηO and a solvent water molecule within the SyLPOR and TeLPOR structures. The well-positioned water, shown in the red sphere, is fixed by the backbone oxygens of Ala91 and Asn115, and the ε-amino group of Lys197. The hydrogen bonds are shown in dashed lines and the bond lengths (Å) are in blue for SyLPOR and dark green for TeLPOR. (B) A proposed proton-relay path following the hydride transfer from NADPH to C17. The photon energy (hv) is represented by a yellow thunderbolt (Dong et al. 2020).

LPOR contains 14 amino acids unique TFT domain that distinguishes LPOR from other structurally related SDR enzymes (Gabruk et al., 2012). The LPOR homologs are structurally conserved with sequence identities of about 54% - 65% between higher plant, cyanobacterial and algal enzymes (Suzuki and Bauer, 1995; Li and Timko, 1996; Dahlin et al., 1999). The secondary structure analysis of LPOR by CD spectroscopy shows 33% alpha-helix, 19% beta-sheets, 20% turn, and 28% random coil (Birve et al., 1996).

### 7.2. Crystal structure of LPOR

Crystal structure of LPORs in their free form (Zhang et al., 2019) and complexed with NADPH have been solved from *Thermosynechococcus elongatus* and *Synechocystis* sp. PCC 6803 at 1.3 Å – 2.4 Å resolution (Zhang et al., 2019; Dong et al., 2020). The above studies highlight the potential importance of hydrogen-bonding networks involving the interaction of LPOR active site residues and Pchlide. The general scaffold of protein remains similar to the typical αβα-topology with a central β-sheet. The crystallographic studies of LPOR demonstrate an 8β-sheet consisting of strands β 3-β 2-β 1-β 4-β 5-β 6-β 7-β 8, the latter being antiparallel. The β-sheets are surrounded by 6 α-helices, (αA, α B, α H) on one side and (αC, α D, α F) on the other (Dong et al., 2020).

The LPOR homologs of *Synecocystis* and *T. elongatus* contain four evolutionarily conserved cysteine residues; Cys38, Cys89, Cys199, and Cys226 around the active site implicated in Pchlide binding and catalysis. Cys-38 and Cys-89 locate at the ends of β2 and β4, respectively, and Cys-199 locates within αF (Silva, 2014; Dong et al., 2020). Cys 226 is in the loop between β6 and αG and is found to be essential for LPOR membrane interaction. During the proton relay pathway through the catalytic tetrad abundant intermolecular polar interactions take place among NADPH, LPOR, and surrounding water molecules with the help of functional groups and backbone atoms to stabilize the cofactor (Dong et al., 2020).

Near the nicotinamide end, a clam-shaped cavity is formed by predominantly hydrophobic and aromatic residues consisting of Leu232, Phe233, His236, Tyr237, Phe240, Phe243, and Phe246 etc (Dong et al., 2020) (Fig 3. Near here). The extra loop of 33 amino acid segments uniquely present in LPOR and absent in other SDR enzyme superfamily members overlaps with certain fragments of the clam–shaped cavity. It participates in Pchlide binding, formation of pigment-complexed POR aggregates and Chlide release (Birve et al., 1996; Reinbothe et al., 2003; Sameer et al., 2021). The LPOR oligomerization takes place upon Pchlide binding which brings about the interaction of the hydrophobic residues and intermolecular interactions in the two distally located lid regions in the POR monomer active site (Gabruk and Mysliwa-Kurdziel, 2015; Zhang et al., 2019, Zhang et al., 2021). It is known that mostly two long α-helices are involved in POR oligomerization (Dong et al., 2020). A POR octamer has been isolated and its structure investigated by cryo-electron microscopy at 7.7 Å resolution. This structure shows that oligomer formation is most likely driven by the interaction of amino acid residues in the highly conserved lid regions (Zhang et al., 2021).

**Figure 3.**
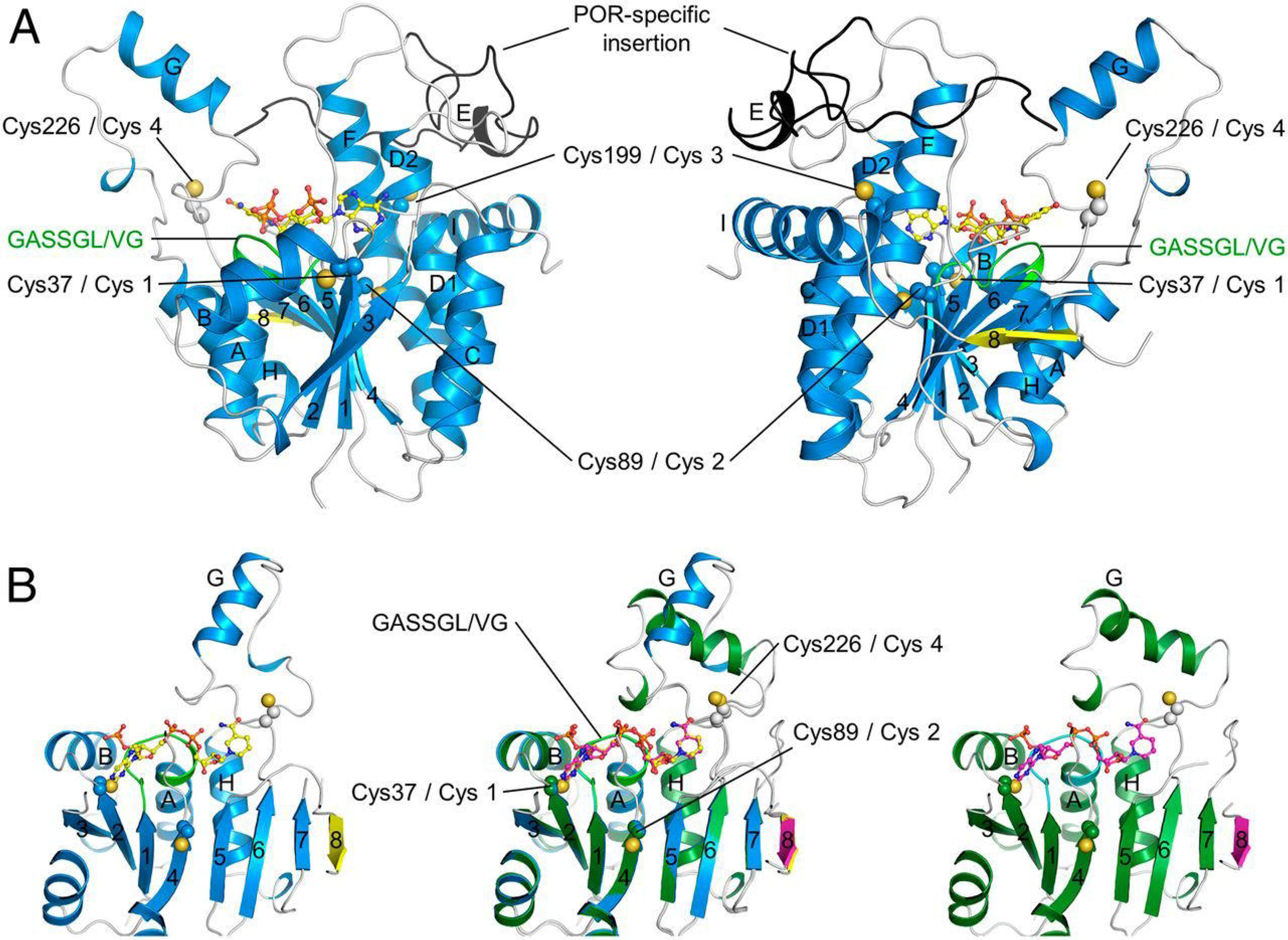
The crystal structure of SyLPOR and TeLPOR from Dong et al. 2020. Ribbon representation of the overall structures of SyLPOR and TeLPOR. (A) Two side views of SyLPOR. The secondary structure elements are colored in blue except the antiparallel β8 in yellow. The loop region is in gray. The LPOR-specific insertion is colored in black. The NADPH-binding sequence is colored in green. Four cysteine residues are shown in sphere mode. The cofactor NADPH is shown in stick-and-ball mode. (B) Front view of SyLPOR (Left), TeLPOR (Right), and their superimposition (Middle). The secondary structure elements of TeLPOR are colored in deep green except β8 in magenta; the NADPH-binding sequence is colored in cyan. The α-helices are labeled alphabetically, and the β-strands are labeled numerically (Dong et al., 2020).

## 8. De-etiolation via LPOR

LPOR was initially isolated and characterized from the etiolated plastids of monocots such as barley, oat, and rye in the prolamellar bodies (PLB) (Apel et al., 1980; Roper et al., 1980; Joyard et al., 1990). It is now known that cyanobacterial LPORs are capable of forming PLBs indicating common features between the cyanobacterial and higher eukaryotic LPOR enzymes (Masuda et al., 2009). When the seed germinates beneath the earth in the absence of light i.e., during skotomorphogenesis, LPOR accumulates to high levels as a macromolecular complex with Pchlide and NADPH in highly organized 3-D lattice-like membrane structure known as prolamellar bodies (PLBs) inside etioplasts (Ryberg and Sundqvist, 1982; Solymosi and Schoefs, 2008; 2010). Both the PLBs and PTs are constituents of the etioplasts which are plastids that have not been exposed to light (Reinbothe et al., 2010).

When etiolated leaves are subjected to a flash of light, the large aggregates of POR–Pchlide– NADPH ternary complexes are converted to POR–Chlide–NADPH complexes. Such ternary complexes have higher emission and are slowly dissociated into smaller complexes accompanied by the progressive release of Chlide from the POR catalytic site. This leads to a large blue shift in absorption and emission maxima of Chlide and is called the Shibata shift. The process ends with the formation of Chlide absorbing at 672 nm and emitting at 682 nm (Chlide682). Crosslinking experiments have shown that Chlide 672-682 is partly composed of Childe still bound to POR complexes and partly by Chlide bound to other proteins (Ryberg et al., 1992; Wiktorsson, 1993). Shibata shift is followed by the formation of photoactive photosystem II (PSII) units (Franck et al., 1999). The Shibata shift is arrested in extreme environmental conditions that impair plastid development (El Hamouri et al., 1981; Dalal and Tripathy, 2012).

During plant development in dark, both the photo- and non-photoactive pools of Pchlide accumulate at different proportions. The Photoreduction of Pchlide to Chlide is mediated by several short lifetime intermediates, e.g., semi-reduced Pchlide radical species formed by hydrogen transfer from NADPH (Belyaeva et al., 1988; Lebedev and Timko, 1999) and characterized by their very low fluorescence yield (Schoefs, 2000). At least three different spectral forms of Pchlide are recognized in intact tissues based on their fluorescence emission maximum (in nm): Pchlide F631, the short-wavelength Pchlide form thought to be located in the tubular lamellae pro-thylakoids bound in a monomeric form to proteins. F644 due to the association of oligomeric LPOR, and Pchlide F655 due to localization in PLBs with polymeric LPOR (Böddi et al., 1992, 1993). The fluorescence lifetime of Pchlide measured in plants showed that short- and long-wavelength Pchlide forms have fast (0.3 to 0.8 ns) and slow (5.1 to 7.1 ns) components with different proportions depending on plant species (Apel et al.,1980; Sundqvist and Dahlin, 1997; Böddi et al., 1998; Kis-Petik et al., 1999; Mysliwa-Kurdziel et al., 2003).

The main photoactive form present in etiolated plants is Pchlide F655, which after illumination is converted to Chlide and subsequently to Chl (F682) through the formation of long wavelength intermediates (Bodd□di and Franck, 1997; Schoefs et al., 2000). The reaction consists of 3 distinct steps including an initial light-driven step followed by dark steps which occur close to or above glass transition temp of proteins. The reduction reaction occurs at temperatures as low as 193K, and in response to femtosecond manipulation of light pulses, signifying its biochemical novelty (Heyes and Hunter, 2005; Heyes et al., 2006). As a result of the light-induced reduction of Pchlide, PLBs disintegrate and the etioplast develops into the chloroplast. The PTs ultimately transform into well-organized thylakoid membranes (Oliver and Griffiths, 1982; Ryberg and Sundqvist, 1988). The isoforms of LPOR are present at different locations of etio-chloroplasts inner membranes (Grzyb et al., 2013; Kowalewska et al., 2016). After flash illumination, the photoactive Pchlide complex can be regenerated by reloading with non-photoactive Pchlide on a fast time scale with concomitant release of Chlide (Franck et al.,1999; Schoefs et al., 2000). Long-term illumination (i.e., greater than a minute) usually converts non-photo-active Pchlide to photo-active Pchlide.

The spectrally different forms of Pchlide in etioplast are attributed to the formation of different size aggregates of LPOR and NADPH. LPORs from different organisms can adopt a variety of soluble oligomeric forms ranging from monomers in cyanobacteria to higher-order oligomers in plant enzymes (Gabruk and Mysliwa-Kurdziel, 2015; Gabruk et al., 2015). Pchlide binding leads to a structural reorganization in the POR enzyme, which in turn induces an interaction of POR monomers (Böddi et al., 1989). POR aggregates account for approximately 95% of total protein in PLBs. LPOR dimerization is not conserved for cyanobacteria species such as SyLPOR and TeLPOR (Dong et al., 2020). In contrast, oligomerization is better conserved in higher plants; species-specific amino acid residues mediate the oligomerization of LPOR. In Arabidopsis thaliana residues 85–88 and 240–270 regions are implicated to participate in oligomerization (Gabruk et al., 2016). Glycerolipids monogalactosyl diacylglycerol (MGSG), digalactosyl diacylglycerol (DGDG) and sulfosyl quinoline diacylglycerol (SQDG) account for a total of 80-20 % of the total lipids in the thylakoids membrane (Solymosi and Schoefs, 2010; Gabruk et al., 2017; Fujii et al., 2017; 2018; Gabruk and Mysliwa-Kurdziel, 2020; Heyes et al., 2021). These lipids play an important role in the aggregation of the Pchlide-LPOR-NADPH complexes, and membrane structure organization in etioplast development (Kobayashi, 2016; Heyes et al., 2021).

There is a possibility of the presence of species-specific motifs in plant LPORs within the oligomerization region. Numerous studies on leaves and isolated plastids indicate that Pchlide: LPOR: NADPH aggregates interact with the lipids of PLB and are responsible for light-triggered PLB dispersion (Engdahl et al., 2001; Gabruk et al., 2017; Fujii et al., 2017; Gabruk and Mysliwa-Kurdziel, 2020). The exact mechanisms for these processes, however, are still elusive (Gabruk and Mysliwa-Kurdziel, 2020). Prokaryotic LPORs from *Gloeobacter violaceus* PCC7421 and *Synechocystis* sp PCC6803 could successfully restore characteristic PLB structures in LPORA knockout mutant of *A. thaliana* even though the size and structure of PLBs were normal, there was a lower ratio of photoactive to non-photoactive Pchlide (Masuda et al., 2009). LPOR overexpression studies in LIPOR deficient cyanobacterium in the dark show the formation of PLB-like ultra-structures in dark. These studies clearly show the intrinsic capability of LPOR to trigger PLBs formation irrespective of its origin in phototrophs (Yamamoto et al., 2020).

## 9. LPOR diversity and phylogeny

LPOR contains multiple isoforms that exhibit differential subcellular localization, expression pattern, mRNA stability, plastid import pathway and response to light. Although POR proteins were known since a long time, the genes coding PORA and PORB were first identified *in A. thaliana* and *H. vulgare* (Reinbothe et al., 1996). Since then, LPOR sequences have been discovered in a number of phototrophs. In higher plant LPOR isoforms show >70% sequence identity for the precursor polypeptides and >80% sequence identity for the mature proteins. The transit peptide region at the N terminal which is not a part of the mature enzyme shows lowest homology (Dong et al., 2020).

In gymnosperms LPOR is encoded by a large multigene family, for instance eleven copies of PORB and two copies of PORA have been identified in (Loblolly pine) *Pinus tadea*, *Pinus mungo, Pinus strobus* (Spano et al., 1992; Forreiter and Apel, 1993, Skinner and Timko, 1998, 1999). The function, mechanism, and localization of different LPOR isoforms have been studied and it varies in different tissues during different developmental stages (Masuda and Takamiya, 2004). *A. thaliana* contains three LPOR isoforms (AtPORA, AtPORB, and AtPORC) (Reinbothe et al., 2010; Sousa et al., 2013; Masuda and Takamiya, 2004; Oosawa et al., 2000; Benli et al., 1991; Armstrong et al., 1995; Su et al., 2001, Pattanayak and Tripathy, 2002). *Zea mays* contains PORA and two PORB orthologs PORB1 and PORB2, latter promoting tocopherol biosynthesis post anthesis. Two POR isoforms are found in *Nicotiana tabacum* (Masuda and Takamiya, 2004), *Lycopersicon esculentum* (Masuda and Takamiya, 2004), *Zea mays* (Horton and Leech, 1975), *Oryza sativa* (Sakuraba et al., 2013; Kwon et al., 2017), *Hordeum vulgare* (Apel et al., 1980; Apel, 1981; Schulz et al., 1989; Holtorf et al., 1995), ornamental plant *Amaranthus tricolor* (Iwamoto et al., 2001) and several other species. A single LPOR gene has been detected in *Synechocystis* sp.strain PCC6803 (Suzuki and Bauer,1995; Fujita et al., 1998; Rowe and Griffiths, 1995; Kaneko et al., 1996), *Plectonema boryanum* (Fujita et al., 1998), *Phormidium lamonosum* (Fujita et al., 1998; Rowe and Griffiths, 1995), *Chlamydomonas reinhardtii* (Li and Timko, 1996), *Marchantia paleacea* (Takio et al., 1998), *Pisum sativum* (Spano et al., 1992), *Triticum aestivum* (Teakle and Griffiths, 1993; Masuda and Takamiya, 2004; Schoefs and Franck, 2003), *Avena sativa* (Darrah et al., 1990; Klement et al., 1999), *Musa* (Coemans et al., 2005) and *Cucumis sativus* (Yoshida et al., 1995;Fusada et al., 2000). PORA is exclusively expressed in etiolated seedlings and its mRNA abundance and its expression declines rapidly upon illumination in *Hordeum vulgare* and several other species (Armstrong et al., 1995; Reinbothe and Reinbothe, 1996; Runge et al., 1996; Masuda et al., 2003; Garrone et al., 2015). PORA is light-sensitive, and it majorly accumulates during skotomorphogenesis and plays a critical role in the etioplast development and photomorphogenesis (Paddock et al., 2012; Gabruk and Mysliwa-Kurdziel, 2015). Overexpression studies of PORA in porB-1 porC-1 double mutant restore the Chl synthesis at varying light intensities indicating that transiently active PORA might be capable of functioning at a range of light intensities (Paddock et al., 2010). In essence, PORA expression is negatively regulated on exposure to light. PORA evolved mostly to protect etiolated seedlings from Pchlide-sensitized singlet oxygen-induced photooxidative damage.

In contrast, PORB transcripts are majorly present in thylakoid membranes in young dark-grown seedlings and in illuminated seedlings. PORB concentration remains unaffected during the change of illumination conditions from dark to light (Lebedev and Timko, 1999; Ha et al., 2017; Buhr et al., 2017). PORB is present right from the seedling development to throughout the life of the plant in mature tissues. PORB closely resembles PORA but there are significant differences between the two enzymes with respect to gene expression, requirements for import of the precursor into the chloroplast ad stability in light. Thus, PORA and PORB have unique functions in etiolated seedlings and at the onset of greening (Aronsson et al., 2000; Masuda et al., 2003; Dahlin et al., 1999, Pattanayak and Tripathy, 2002, 2011).

PORC mostly evolved in Brassicales is additionally light indulgent than either PORA or PORB. PORC is expressed in a light intensity dependent manner, being highly expressed in high light (Oosawa et al., 2000; Su et al., 2001; Pattanayak and Tripathy, 2002). PORC mRNA accumulates only after illumination in etiolated seedlings and is predominantly detected in fully matured green tissues during development and throughout the life of the plant (Su et al., 2001; Pattanayak and Tripathy 2002, 2011; Paddock et al., 2010). Despite the physiological equivalence and a perceived redundancy in PORB and PORC functions in mature plants under normal growth conditions it is seen that PORC is differentially regulated and is not under circadian control like PORB. The PORC transcripts are positively regulated by increasing intensity of light while PORB mRNA decreased partially under high light conditions in *Arabidopsis*. Thus, PORB although constitutively active from the seedling stage to the mature plants it is less active under high light conditions (Masuda et al., 2002).

Based on the biochemical analysis and evolutionary studies Gabruk and Mysliwa-Kurdziel 2020, proposed two group of LPOR enzymes- a) Z type LPOR - bacterial origin and b) Plant origin LPOR-S type (AtPORC type active enzymatically active without lipids) and L type LPOR (are active when bound to lipid membrane).

### 9.1. Phylogeny of LPOR isoforms

In the present study we analysed 270 LPOR protein sequences of AABPs, cyanobacteria, algae, bryophytes, pteridophytes, gymnosperms and angiosperms along with LPOR sequences from non-photosynthetic organisms having the characteristic SDR catalytic motif YxxxK and the conserved sequence (Gly-X-X-X-Gly-X-Gly) from the publicly available databases. A Maximum likelihood phylogenetic tree was constructed with Mega 7 software after aligning the sequences with MUSCLE multi-sequence alignment program (Kumar et al., 2016). The phylogenetic distribution was inferred using the boot strap method (500 replicates) with WAG (Whelan and Goldman matrix) and Freq (+F) model. Similar results were obtained when the above LPOR sequences were analysed by neighbor-joining method and UPMGA (Unweighted pair group method with arithmetic mean) (Whelan and Goldman, 2001) (Figure 4. Near here).

**Figure 4.**
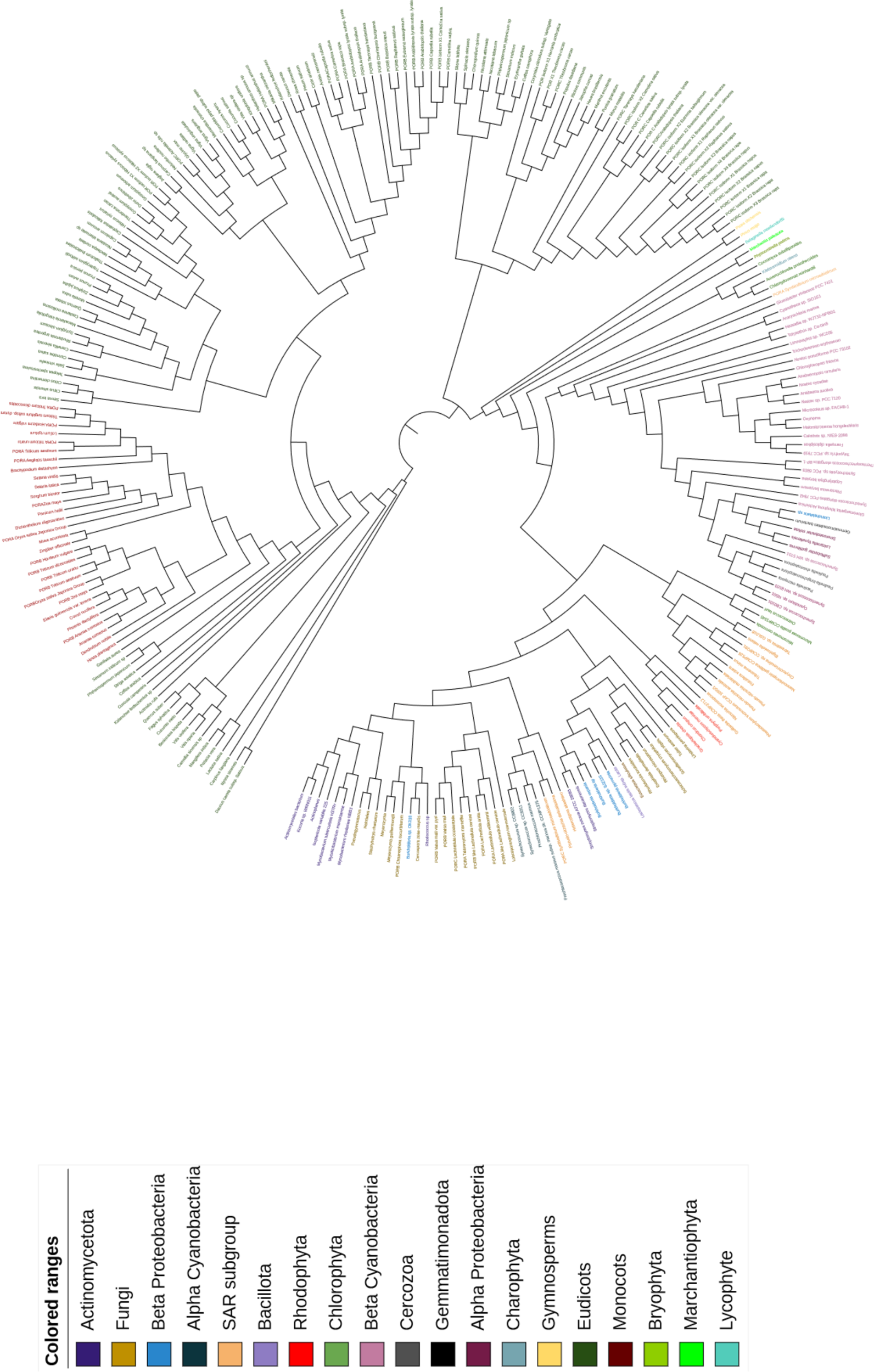
Molecular Phylogenetic analysis by Maximum Likelihood method The evolutionary history was inferred by using the Maximum Likelihood method based on the Whelan and Goldman + Freq. model (Whelan and Goldman, 2001). The bootstrap consensus tree inferred from 500 replicates (Felsenstein,1985) is taken to represent the evolutionary history of the taxa analyzed (Felsenstein,1985). Branches corresponding to partitions reproduced in less than 50% bootstrap replicates are collapsed. The percentage of replicate trees in which the associated taxa clustered together in the bootstrap test (500 replicates) are shown next to the branches (Felsenstein,1985). Initial tree(s) for the heuristic search were obtained automatically by applying Neighbor-Join and BioNJ algorithms to a matrix of pairwise distances estimated using a JTT model, and then selecting the topology with superior log likelihood value. The analysis involved 270 amino acid sequences. All positions with less than 95% site coverage were eliminated. That is, fewer than 5% alignment gaps, missing data, and ambiguous bases were allowed at any position. There were a total of 282 positions in the final dataset. Evolutionary analyses were conducted in MEGA7 (Kumar et al., 2015). A midpoint rooted circular phylogenetic tree was created and annotated in Interactive Tree of Life (Letunic and Bork, 2016; iTOL, https://itol.embl.de/).

LPOR sequences in the present study have been split into 4 major clades, clade 1 shows a deep branching radiation representing LPOR sequences from phylum actinomycetota order Actinomycetales, Micrococcales, Mycobacteriales) and fungal division Ascomycota (orders Incertae sedis, Leotiomycetes sodariomycetes, Saccharomycetales, Mucorales, Helotiales, Eurotiales, Mortierellales) interspersed with beta proteobacteria Burkholderia, α-cyanobacteria and SAR (Stramenopiles-Heterokonts-Alveolates) supergroup*. Streptomyces sviceus*, *S. davaonensis*, *Lactococcus latctis* subsp. lactis lie close to Burkholderiales. Phaeophycean *Ectocarpus siliculous* remains close to beta proteobacterial sequences and fungal LPOR sequences.

There is a clear phylogenetic clustering of LPOR sequences from Sporidiobolales, Chaetothyriales, Mucorales, Schizosaccharomycetales, Saccaromycetales, Pezizales, and Kickxellaleslies closer to beta proteobacterial sequences. Thus, members of Actinomycetota, Psuedomonadota lie close forming a distinct group. The anaerobic and facultative anaerobic non photosynthetic bacteria show diversification of LPOR enabled by HGT.

LPOR in AAPBs is involved in photoconversion of Bacterioprotochlorophyllide to Bacteriochlorophyllide. LPOR sequences from Gemmatimonadetes, betaproteobacteria Limnohabitans, alphaproteobacteria -*Sulfitobater*, *Loktanella* and *Dinoroseobater* lie close to β-cyanobacterial sequences. PORC sequence from algae *Symbiodinium microadriaticum* and LPOR sequences from chlorophyta, charophyta, bryophytes, marchnatiaophyta, lycophyta and gymnosperms cluster together closely with a high bootstrap percentage. Consistent with the previous studies the LPOR of *Gemmatimonadetes* bacterium clusters within beta proteobacterial LPORs as a sister of Limnohabitans.

Presence of LPOR sequences in fungal and other non-photosynthetic bacteria may be a result of infrequent HGT from photoautotrophs to heterotrophs including bacterial and fungal pathogens. *Choanephora. cucurbitarium* LPOR sequence is in all likelihood transferred from prokaryotic or eukaryotic phototroph to fungi as a result of overlapping ecological niche or pathogenic close association (Richards et al., 2009; Tunjić and Korač, 2013; Qiu et al., 2016; Min et al., 2017). LPOR sequences might be involved in Mycobacterium and fungi in certain reduction reactions.

Clade 2 majorly includes many sequences from β-cyanobacteria, algae and certain anoxygenic photosynthetic bacteria. The β-cyanobacterial clade remains separated from α cyanobacterial sequences. SAR LPOR sequences from Gillardia, Cyanidioschyzon merolae, Chondrus crispus, Gracilariopsis chorda, Porphyra umbilicalis, Symbiodinium microadriaticum PORA form a distinct group with 98-60 % bootstrap percentage.

The placement of AAPBs close to picocyanobacteria suggests these LPOR sequences have been acquired by AAPBs through HGT from oxygenic phototrophs (Gabruk and Mysliwa-Kurdziel, 2020: Chernomor et al., 2021). LPOR sequences were transferred from picocyanobacteria to α proteobacteria and then transferred to beta proteobacteria (Chernomor et al., 2021). The LPOR of Gemmatimonadetes cluster with Limnohabitans suggesting a HGT from Limnohabitans to Gemmatimonadetes.

Members of cyanobacteria-Gloeobacter, Cyanothece, Hassalia, Tolypothrix, Limnorphais, Nostoc, Chlorogloeopsis, Anabaena, Microcoleus, Oxynema, Halomicronema, Calothrix, Fremyella diplosiphon, Thermosynechoccus elongates, Synechocystis, Plectonema boryana, Synechococcus, Cyanobium form a separate group in clade 2 with 76% bootstrap percentage. These cyanobacterial species might have evolved LPOR in a certain environmental niche habitat (Pattanaik et al., 2011).

The recent endosymbiont *Paulinella chromatophora* that engulfed alpha cyanobacterium retained the alpha cyanobacterial LIPOR in its chromatophore (Marin et al., 2007; Kim and Park, 2016; Vedalankar and Tripathy, 2019). However, its nuclear LPOR sequence is closer to β cyanobacteria *Synechococcus sp. WH5701*, *Cyanobium*, *Plectonema boryanum* and other species. The presence of LPOR in *P. chromatophora* is a result of HGT from β cyanobacteria suggesting a polyphyletic origin of POR in *Paulinella* and possibly in several other organisms. The Paulinella chromatophora LPOR sequence closest neighbour are *Marchantia palea* and *Phsycomitrella* are positioned on the same branch and their closest neighbour is *Selaginella*. The PORA and PORB sequences are positioned close to each other but on separate branches. Amongst angiosperm PORB is closer to PORC sequences.

The chromatophore genome is highly reduced, encoding 867proteins that represent about one-third of proteins of its free-living counterparts (Nowack et al., 2008). Similar to the EGTs found in Archaeplastida, *P. chromatophora* has relocated> 70 chromatophore genes into the nuclear genome (mostly involved in photosynthesis-related functions) (Nowack et al., 2016; Zhang et al., 2017). By contrast, these genes represent<1% of the *Paulinella* nuclear genome, while in *A. thaliana* some reports have suggested that the genes of cyanobacterial origin can account for upto 18% of the nuclear genes (Martin et al., 2002).

The clade 3 and 4 mostly includes eukaryotic LPOR sequences from angiosperms consistent with the other LPOR phylogenetic studies. The angiospermic LPOR sequences are distinctly divided into–eudicots and monocots. Monocots members from Poales, Arecales, Zingiberales and asparagales lie close together. *Musa acuminate* LPOR lies close to *Zingiber officinale* with a 62% bootstrap percentage. Poales from a distinct group with 88% bootstrap percentage. All the monocot sequences are placed in between two distinct dicot groups.

Eudicots group - Brassicales, rosales, myrtales, malpighiales, malvales, gentianales, lamiales, solanales, caryophyllales, fabales, rosales, apiales, saxifragales, asterales, ranunculales, vitales, cucurbitales, ericales, fagales, malvales, oxalidales, celastrales, proteales, sapindales lie close together forming one separate group of eudicots. Certain members of lamiales, gentianales, solanales, saxifragales, ericales, fagales, cucurbitales, sapindales, asterales, cornales and apiales from the second group of eudicots. PORA and PORB sequences are clustered together whereas PORC sequence form a completely distinct subgroup among eudicots suggesting a completely different selection pressure that led to their evolution.

Most of the sequences analysed distinctly show that despite a great deal of similarity between the LPOR isoforms. These isoforms form separate nodes with PORC present at a distance from PORA and PORB suggesting that PORA and PORB must have evolved around the same time and PORC evolved later as a result of the selection pressure. In this context it is to note that the wild diploid Brassicales and related hybrids evolved in inhospitable conditions with abilities to withstand drought, heat, high light and salt stresses (Arias and Pires 2012; Folk et al., 2020). This explains the evolution of LPOR in Brassicales in PORC gene is usually expressed in high light conditions (Pattanayak and Tripathy, 2011) to protect plants from ROS mediated oxidative stress.

The duplication events gave rise to LPOR isoforms and independent duplication events happened in monocots and dicots. The LPOR gene kept duplicating with the diversification of angiosperm species. Both primary and secondary duplication are responsible for the spread of LPOR. HGT and EGT have played an exceedingly huge role in forging the evolution of LPOR in phylogenetically diverse lineages (Gabruk and Mysliwa-Kurdziel, 2020).

## 10. Conclusion

Evolutionary studies validate the premise that cyanobacterial and plant LPORs originated in the oxygenic Earth about 1.36 bya from a single common ancestor. LPORs originated in the oxygenic Earth about 1.36 bya from a single common ancestor. The phylogenetic studies clearly show a deep branching pattern in LPOR with a great degree of similarity between the PORA and PORB in different taxa suggesting a closer relationship between them. PORC originated in Brassicales in a separate duplication event as a result of unfavourable climatic conditions (Gabruk and Mysliwa-Kurdziel, 2020) to protect plants from environmental stresses.

LPOR is a more evolved Pchide reducing enzyme and loss of LIPOR gene from several eukaryotic lineages clearly show the importance of LPOR in synthesis of chlorophyll, phytochrome mediated developmental processes and protection of plants from ROS induced oxidative stress. Climatic conditions of increasing oxygen tension in atmosphere, high light, high temp, drought, heat along with nutrient status led to multiple gene losses and gene duplication events that played a big role in the current distribution of LIPOR and LPOR. The evolution of LPOR paved way for the loss of LIPOR from highly evolved photosynthetic organisms. The exclusive inheritance of LPOR in angiosperms is a result of differential plastidic LIPOR gene loss. The coexistence of LIPOR and LPOR ensures that there is a continuous supply of chlorophyll under altered (light, oxygen, temperature and other environmental conditions.

It would be a challenge to determine if LIPOR and LPOR originated from a common ancestral prokaryote. The distribution of LPOR is both a result of chance and contingency and the relationship between LPOR and LIPOR seem to be less straightforward when looked through the analysis of a few taxa. Our phylogenetic analysis indicates that LPOR is present in a wide set of photosynthetic (AABPs, Cyanobacteria, algae, bryophytes, pteridophytes, gymnosperms, angiosperms) and non-photosynthetic (bacteria and fungi) organisms. LPOR sequences from non-photosynthetic organisms (actinomycetota, bacillota, ascomycota, basidiomycota, mucoromycota, Zygomycota, zoopagomycota) lie close to prokaryotic LPOR sequences. Although no potential functional significance can be assigned to these LPOR, these might contribute to some other specific physiological role in the lifestyle of the organism. Phylogenetic incongruence indicates that LPOR genes have been overwhelmingly horizontally transferred between not only photosynthetic but also non photosynthetic lineages.

In this context it would be interesting to see if the signature SDR motif in the fungi is coding for functions yet to be understood. The identification of the function of these LPOR sequences would give us a clue to the LPOR migration and evolution. Many more cases of intra and interphylum HGT events that are yet to be discovered will shed better light on the evolution of protochlorophyllide oxidoreductase.

## Acknowledgment

We are thankful to the authors and the journal, Proceedings of National Academy of Sciences, USA for allowing us to use crystallographic structure of cyanobacterial light-dependent protochlorophyllide oxidoreductase (Dong CS, Zhang WL, Wang Q, Li YS, Wang X, Zhang M and Liu L (2020) Crystal structures of cyanobacterial light-dependent protochlorophyllide oxidoreductase. Proceedings of the National Academy of Sciences 117(15):8455-8461.)

## Abbreviations

endosymbiotic gene transfer: EGT
horizontal gene transfer: HGT
light-dependent protochlorophyllide oxidoreductase: LPOR
light-independent protochlorophyllide reductase: LIPOR

